# Evolutionarily Optimal Phage Life-History Traits: Burst Size vs. Lysis Time

**DOI:** 10.1101/2025.04.16.649164

**Authors:** Joan Roughgarden

**Affiliations:** Hawaii Institute of Marine Biology, University of Hawaii, Department of Biology, Stanford University

## Abstract

A new model based on a dynamical equation for the virus to microbe ratio (VMR) during log phase population growth shows that an optimal balance occurs between a short lysis time with low burst size *vs*. a long lysis time with large burst size. The model predicts that interventions lowering phage adsorption by killing free virus and/or limiting their access to bacteria favors the evolution of an increased lysis time and higher burst per infecting microbe until the intervention either drives a virulent phage extinct or, for temperate phage, drives the phage from its lytic phase into its lysogenic phase. The model also predicts that along an environmental gradient of increasing primary productivity the optimal lysis time shortens along the gradient, implying that the lytic life cycle goes around faster along the gradient.

**Importance:** A new approach to modeling phage life history predicts that virus respond to interventions that limit their adsorption onto bacteria by evolving a longer lysis time. The new model also predicts that lysis time of virus in nature shortens and the virus life cycle goes around faster as environmental conditions favoring virus production increase. These predictions show that virus life-history traits are not arbitrary and can be predicted in advance based on environmental conditions.

## Introduction

Breitbart (2012) noted that “It is difficult to find a paper in marine viral ecology that does not begin with a statement about the sheer abundance of viruses”. The same is true today. The number of prokaryotes on earth is 4–6 x 10^30^ cells with a production rate of 1.7 x 10^30^ cells/yr (Whitman *et al*. 1998). The number of virus is about ten times the number of prokaryotes (Wommack and Colwell 2000) implying that the number of virus is on the order of 10^31^. In comparison, the number of grains of sand on all the earth’s beaches is only 4 x 10^20^ and the number of stars in the universe a meager 2 x 10^19^ (Plait 2025).

Furthermore, virus are ancient with differences so great among them that the taxonomic rank of “realm” has been coined to reside at atop 15 lower ranks that include multiple kingdoms as well as the taxa below that (ICTV 2020, Koonin *et al*. 2020, 2021, Harris and Hill 2021). Marine phage represent 94% of the nucleic-acid-containing particles in the oceans (Breitbart *et al*. 2018). Hence, phage represent an enormous diversity of evolutionary agents whose phenotypes are shaped by natural selection.

What then are the phage phenotypes that natural selection is shaping? Of the many traits phage possess such as their morphology and mechanisms of inheritance and replication, their life-history traits are of particular interest. These traits include the lytic/lysogenic switch, the burst size and latency.

Conditions for the switch between lytic and lysogenic life cycles were developed in a previous paper (Roughgarden 2024). Predictions from that paper’s model can account for eco-geographic patterns of the ratio of lytic to lysogenic phases.

This paper focusses on the tradeoff between burst size and latency. However, information on possible eco-geographic patterns in burst size and latency across habitats is limited. Both traits vary considerably across and within habitats, often in accordance with transient environmental conditions such as temperature (Wommack and Colwell 2000, Weinbauer 2004, Greenrod *et al*. 2024). Parada *et al*. (2006) show a correlation between burst size and bacterial productivity in coastal waters, but do not present data on latency times. Ranasinghe (2019) offers an extended review of burst size and latency across various categories of virus, but does not report the habitats where the virus live. This article extends the model in (Roughgarden 2024) to explain burst size and latency and predicts possible eco-geographic patterns whose discovery would test the model.

Interventions to control microbes with antibiotics have produced antibiotic resistance (Martinez *et al*. 2008, Davies and Davies 2010). These and other interventions might affect virus as well. The model in this article also predicts the evolutionary consequences for phage of interventions that lower the overall adsorption rate by virus onto bacteria.

This article explains burst size and latency as an evolutionarily optimal balance between reproducing early to benefit from geometric growth versus reproducing late to benefit from the time to manufacture a large burst size.

The study of an evolutionary tradeoff between the benefits of early and late lysis time enjoys a long history including, most recently, Wang 2006, Bull 2006, Heineman and Bull 2007, and Kannoly *et al*. 2022. The models developed in this literature present a system of delay-differential equations. Mass action virus/bacteria collisions describe the process of viral infection. The bacteria population grows with density dependence. These equations are solved for the equilibrium abundance of uninfected bacteria, infected bacteria and free phage.

This article differs from the preceding literature in being based on a new model defined in discrete time. It features viral infection of bacteria as a Poisson process and assumes density dependence in viral adsorption to bacterial surfaces. The model pertains to a virus-bacteria system in the log phase, where both populations grow exponentially rather than exist at equilibrium. The model anticipates a bust-boom environment where bacteria frequently experience exponential growth rather than a stable environment where an equilibrium can be attained.

Thus, this paper envisions the situation in which the optimal life history evolves differently compared to the earlier literature—log phase versus stationary phase, recursion equations in discrete time versus delay-differential equations in continuous time, Poisson infection versus mass action infection and density dependence in the virus rather than in the bacteria. The section where the model is introduced discusses the reasons why a new model was needed. The Discussion section features a comparison of the differing intuition behind predictions from the earlier models compared to the present model.

Two studies (Wang 2006, Kannoly *et al*. 2022) demonstrate experimentally that an optimal balance between lysis time and burst size exists. Hence, the reality of an optimal balance can now be taken as established. Instead, this article concerns how to predict *what* that optimal balance is. In nature, phage may not always possess the optimal balance for their present circumstances, but knowing the optimal balance in those circumstances reveals the direction toward which evolutionary change will occur.

The article has five sections: The first describes the basic population-dynamic model. The second extends the model to predict the optimal lysis time and burst size. The third compares the predicted the optimal lysis time and burst size to data from experiments with *λ*-phage and *Escherichia coli* (Wang 2006). The fourth predicts the evolutionary impact of interventions that affect the adsorption rate. The fifth develops predictions for eco-geographic patterns in lysis time and burst size.

## Model

### Population Dynamics of Phage and Bacteria

#### The Classic Model

One might wonder why a model of phage/bacteria population dynamics is being used that differs from the traditional model and its many extensions originally introduced by Campbell (1961). Here is why.

The Campbell model is an adaptation of the standard Volterra predator/prey model in ecology (Volterra 1926, *cf*. Roughgarden 1998):

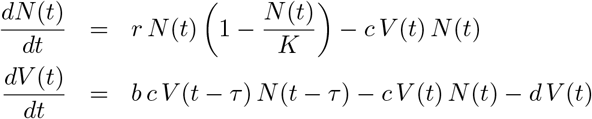

where *N* (*t*) is the number of bacteria at time *t, V* (*t*) is the number of virus particles at time *t, N* (*t* − *τ* ) is the number of bacteria at time *t* − *τ, V* (*t* − *τ* ) is the number of virus particles at time *t* − *τ, τ* is a time lag, *r* is the bacterial intrinsic rate of increase, *K* is the bacterial carrying capacity within its environment, *c* is the rate of conversion of mass-action collisions between virus particles and bacteria into infections, *b* is the burst size, and *d* is the per capita disintegration rate of virus particles. (The Campbell model was originally written for a phage/bacteria system in a chemostat. Here the chemostat nutrient flow terms are omitted. Also, both *N* (*t*) and *V* (*t*) are numbers, not densities. The equations for the numbers cannot be converted into equations for densities by dividing both sides with the volume of a vessel—the mass action term is a product of numbers that cannot be converted into a product of densities.)

The Campbell model is a pair of nonlinear delay-differential equations. The first term of the equation for the bacteria assumes they grow logistically in the absence of virus. The second term is the loss of bacteria from infection brought about by mass-action random collision of bacteria with virus particles. The first term in the equation for the virus assumes the production of new virus particles depends on the collisions that took place *τ* units of time previously. The time lag, *τ*, called the latency, is the time needed for the infecting virus particles to manufacture and release new virus particles using the bacteria’s replication machinery. The next term indicates the loss of free virus particles from the infections taking place at the present time, *t*. Thus, the virus particles disappear for the duration of the latency period. After infecting bacteria at time *t* − *τ* they reappear at time *t* as newly minted virus particles. And the last term is the natural decay of virus particles.

The equilibrium bacterial abundance, 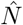, is 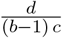 independent of *K*, whereas the equilibrium viral abundance, 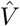, is 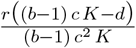, which does depend on *K*. Indeed, *K* must exceed 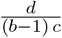 for 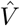 to be positive. Thus, the viral abundance at equilibrium depends on the bacterial carrying capacity whereas the bacterial abundance does not depend on its own carrying capacity because its abundance is controlled by the virus. The equilibrium is a stable focus for *τ* = 0 but is unstable for any *τ >* 0 (*cf*. Roughgarden 2024, Supplementary Material).

Krysiak-Baltyn *et al*. (2017) published an extensive review of models for virulent phage and bacteria. They observed that “The basic model initially suggested by Campbell, and later refined by Levin and colleagues [Levin *et al*. 1977], has remained in use since first publication. Later studies have mostly made minor adjustments to the existing model to study dynamic behavior.” (p. 964) They further observed that “Earlier studies on simple chemostat models indicated that the dynamics of the model are not in good quantitative agreement with experimental data. In particular, the stability of the simulated models is lower than expected, with many exhibiting heavy oscillations and extinction events where stable steady-states exist experimentally.” (p. 964) Krysiak-Baltyn *et al*. (2017) conclude that “On a practical level, the usefulness of computational models of bacteria and phages has not yet been established. The simpler chemostat models may first need to become more accurate” (p. 965).

The Campbell model has quantitative limitations. First, it predicts instabilities to virulent virus/bacteria population dynamics that are contradicted experimentally. Second, the Campbell model is so mathematically challenging that its use is impractical in biology. How to mathematically analyze a system of nonlinear delay-differential equations is an ongoing subject of research in applied mathematics (Tang and Zou 2008, Huang *et al*. 2010; indeed many investigators simply put the time lag, *τ*, equal to zero to evade the problem, converting the delay differential equations into ordinary differential equations).

The Campbell model is also qualitatively questionable. The Campbell model envisions that virus-/bacteria population dynamics is based on two counteracting forces—density-dependence in the bacteria that counteracts the Volterra model’s predator/prey oscillations, exacerbated by time lags. The outcome is a balance of these forces. Is this qualitative picture what is actually going on in the phage/bacteria interaction? This paper argues otherwise.

Moreover, the formulas for the virus/bacteria equilibrium and its stability depend both on properties of the microbial population dynamics as well as on parameters for the virus/bacteria interaction. This commingles parameters for the virus/bacteria interaction with those for bacterial population growth.

Commingling virus/bacteria interaction parameters with parameters for bacterial population growth leads to peculiar predictions. For example, according to the formula for 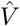 in the Campbell model, the bacteria in the guts of small mammals should have fewer strains of phage infecting them than bacteria in large mammals because the low bacterial carrying capacity in small guts precludes the coexistence of many types of phage. Conversely, the bacteria in the guts of large mammals should support more strains of phage because the large carrying capacity for bacteria in large guts allows more types of phage to coexist with bacteria. But, according to the Campbell model, any increased phage/bacteria coexistence resulting from a large bacterial *K* in large-mammal guts is temporary because the weakened density dependence from a large bacterial *K* leads to large virus/bacteria oscillations. None of these predictions has been observed.

For all these reasons, this paper views the Campbell model and its extensions as an unstable foundation for deriving optimal life-history traits. Instead, an alternative model of phage/bacteria population dynamics from (Roughgarden 2024) is used.

#### Alternative Model

Although the alternative model here avoids the specific difficulties of the Campbell model, it introduces a different set of simplifying assumptions that themselves might be questioned— there is no perfect model.

The model here assumes no density dependence in bacterial population growth. The stability of virus/bacteria coexistence is determined solely by properties of the virus/bacteria interaction itself as determined by functions describing how burst size relates to the multiplicity of infection and how the adsorption of virus onto bacterial surfaces relates to the virus to microbe ratio.

The model pertains to the log phase of bacterial population growth. Once a stable virus/bacteria equilibrium *ratio* is reached during the log phase, both parties possess a common geometric growth factor. Thereafter, external to the virus/bacteria interaction, density dependence might enter the bacterial population dynamics. If so, after attaining coexistence at a stable virus/bacteria ratio during their log phase, the population pair can grow together eventually reaching a joint stationary phase. The focus in this paper however, is not on the stationary phase—this was analyzed in the prior paper, but on conditions for the initial colonization by virus into an uninfected bacterial population.

The model is defined in discrete time, where the time step is the lysis time. Virus particles are produced anew each step. Any virus particles not adsorbed are lost or become inactive. The burst of virions produced during each time step is analogized to pollen. Oak trees fill the air with trillions of pollen grains some of which adsorb to stigma and the rest disappear. The next year the cycle repeats. The lysis time is the time step in this model analogous to how the generation time is the time step in population-genetic models.

The kinetics of infection in the model consists of Poisson sampling from an environmental pool of virus particles instead of mass action collisions. Poisson infection of bacteria by phage has been well documented long ago (Ellis and Delbrück 1939). Accordingly, after the infection process, some bacteria remain uninfected, some have only one infection, some have two infections *etc*. Poisson searching for hosts also features in early parasitoid/host models (Nicholson and Bailey 1935) and more recently in models for bacterial colonization of hosts (Roughgarden *et al*. 2018, Roughgarden 2020, 2023).

Readers have expressed misgivings about three features of this model. First is the assumption that unadsorbed virus are unavailable for subsequent time steps. The lysis time for phage can be from half an hour to several hours (Kutter and Sulakvelidze 2004). The longevity of phage in the environment can be days to years (Jończyk *et al*. 2011). It is thus possible that phage produced at one time step can infect at later time steps. The issue however, is not longevity relative to lysis time but whether phage from previous time steps remain in the environmental source pool supplying the current infections. The phage from previous time steps can be disbursed and diluted through wind, currents, exposed to UV radiation and so forth. The simplifying assumption here is that unadsorbed phage from preceding time steps are rare and practically unavailable to the viral source pool for current infections. If carry-over infection is important, then a future extension of the model could treat long-lived virus in the environment analogous to the seed bank in plant population models (*eg*. Jarry *et al*. 1995, Eager *et al*. 2013).

A second feature drawing misgivings is the absence of density dependence in the bacteria. Many experimental studies of the phage/bacteria interaction take place in chemostats and other vessels with fixed volume in which the assumption of bacterial density dependence is natural and realistic. The intuition for the model here however, pertains to virus and bacteria in a natural habitat like soil and plankton subject to bust-boom environmental shocks. In this situation, the bacterial population is rarely abundant enough to incur density effects and instead is continually in a state of recovering from the most recent climate event. If density dependence is important, then a future extension of the model could add a bacterial carrying capacity parameter and the formulas reworked accordingly.

A third feature of concern has been use of the word, “extinction”. Here, extinction means that the *ratio* of virus to microbe tends to zero, not that the virus population itself tends to zero. Because both populations are growing (or declining) exponentially, their ratio is the appropriate definition of extinction here.

These qualifications notwithstanding, the premise of this paper is that sufficient detail about the phage/bacterial interaction has been included to support predictions concerning viral life-history evolution.

Turning now to how the model is formulated, Figure 1 shows a lytic life cycle on the left and a lysogenic life cycle on the right. A time step begins with the state variables, *N* (*t*) and *V* (*t*) for the number of bacteria and free virus particles (virions), respectively, in their environmental source pools at time *t*. The *v*(*t*) ≡ *V* (*t*)*/N* (*t*) is the ratio of virus to bacteria in their source pools (the environmental virus to microbe ratio, VMR).

**Figure 1:**
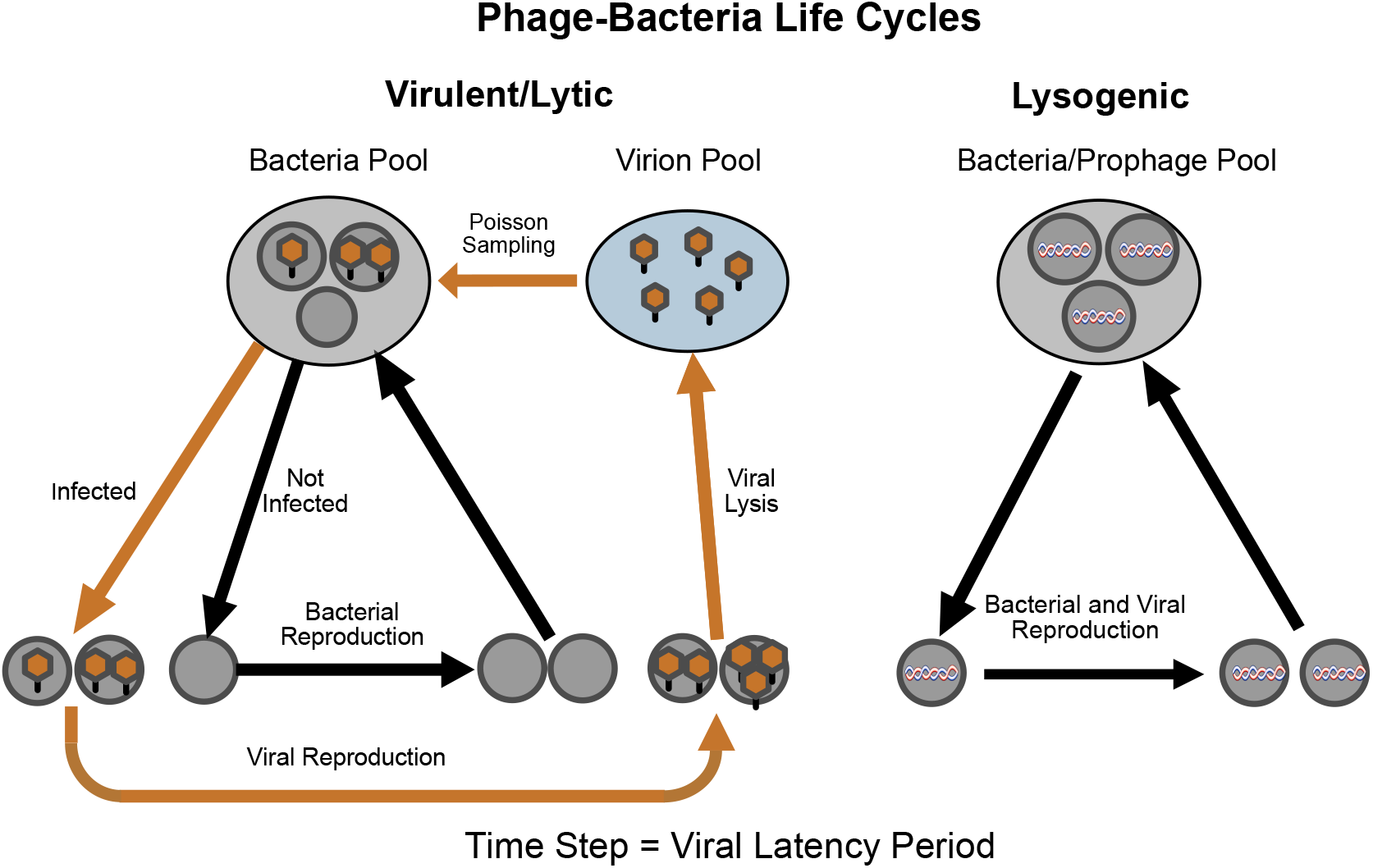
Phage/bacteria life cycles. Left: life cycle for virulent phages and for the lytic phase of temperate phages. Bacteria in the bacterial source pool in the environment are infected by free virus particles (virions) from the viral source pool according Poisson sampling. The infected bacteria are the site of viral replication, leading to the release of new virions and the death of the infected cells. The uninfected bacteria replicate to produce more bacteria available for infection to begin the next time step. Right: life cycle for the lysogenic phase of temperate phages. The virus is represented as a helix that is incorporated as a prophage into the genome of its bacterial host. The virus life cycle is coincident with the bacterial life cycle. The time step is the viral latency period.

The virus infect the bacteria with Poisson sampling such that some bacteria remain uninfected, some have only one infection, some have two infections *etc*. The Poisson distribution describing the infection process has density parameter, *µ*(*t*) ≡ *a*(*v*(*t*)) *v*(*t*). The *a*(*v*(*t*)) is the adsorption function— it refers to the process whereby a viral particle comes in contact with a bacterium, binds to surface molecules, and then is incorporated into the bacterial cell. Adsorption is a function of *v*(*t*) because the number binding sites on the surface of the bacterium is finite and the sites may become saturated.

The *a*(*v*(*t*)) summarizes information about the binding affinity of virus particles to receptors in the bacterial surface, the number of available binding sites and the accessibility of the bacteria to virus particles. The accessibility is influenced by how dense the bacteria are as well as by how viscous the media is, or for airborne virus particles, whether air filters are installed.

With a Poisson distribution, *µ* is the average total adsorption per time step for a given *v*. Upon measuring *µ*, the adsorption coefficient is calculated as *a* = *µ/v* for the given *v*. Repetition for several values of *v* yields several values of *µ* from which the corresponding values of *a* are calculated. The values of *a* are then fitted to a curve yielding the adsorption function, *a*(*v*). A detailed illustration is noted below.

According to a Poisson density function, the fraction of bacteria that remain uninfected, *P*_0_(*µ*(*t*)), is *e*^−*µ*(*t*)^ and the fraction of bacteria colonized by *m* virus, *P*_*m*_(*µ*(*t*)), is 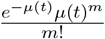 for *m* ≥ 0. The *m* indexes the number of virus per bacterium, called the multiplicity of infection (MOI). The fraction of bacteria that are infected by one or more virus is 1 − *P*_0_(*µ*(*t*)).

The uninfected bacterial population increases by a geometric factor of increase per time step, *R*. If *R >* 1, the bacteria can increase in the absence of the virus. Accordingly, the uninfected bacteria resupply the bacterial source pool as

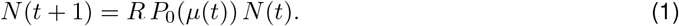

Thus, the population size of bacteria at time *t* + 1 equals *R* times the number of bacteria that are not infected. The *R* refers to microbial population growth over the time step.

Meanwhile, the virus within the infected bacteria produce new virus particles as

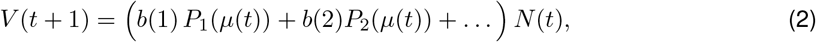

where *b*(*m*) is the burst size from a bacterium infected with *m* virus particles. Thus, the population size of virions at time *t* + 1 equals the number of virions produced from bacteria infected by one phage, plus those produced from bacteria infected by two phage, plus those from bacteria infected with three phage, and so forth. No unadsorbed phage carry over to the next time step—they are lost. The phage at each time step are newly produced as bursts from infected bacteria. Together, Eqs. 1 and 2 constitute a dynamical system consisting of coupled recursion equations for bacteria and phage.

The viral adsorption depends linearly on the virus to microbe ratio, *v*,

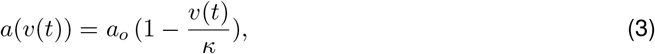

where *κ* is the saturation coefficient—the virus to microbe ratio at which all the binding sites per bacterium are occupied and *a*_*o*_ is the maximum adsorption coefficient that occurs when all the binding sites are available.

To illustrate how to measure the adsorption function, suppose a vessel of known volume is provided with 10,000 phage and 100 bacteria—this corresponds to *v* = 100. After a time step, determine how many phage are left. Say, the number phage left in one replicate is 9,990 indicating that 10 phage were adsorbed. In two more replicates, the adsorbed number was 12 and 8, respectively, so the average is 10 adsorbed—this is *µ* for *v* = 100. Therefore, *a*(100) = *µ/v* = 10*/*100 = 0.10. Suppose further that this experiment is repeated for various *v*. If the vessel is provided with 20,000 phage and 100 bacteria, corresponding to *v* = 200, then say, 14 are adsorbed on average. Therefore, *a*(200) = 14*/*200 = 0.07. Continuing, with 30,000 phage and 100 bacteria, *v* = 300, the number adsorbed turns out to be 15 on average so that *a*(300) = 15*/*300 = 0.05. Thus, total adsorption, *µ*, increases with *v*, while the adsorption per unit VMR, *a*, decreases with *v* as the receptor sites fill up. The best fit of a line to the points, (100, 0.10), (200, 0.07), (300, 0.05), is *a*(*v*) = 0.12333 − 0.00025 *v*. The Y-intercept of the line is *a*_*o*_ = 0.12333, the maximum adsorption coefficient pertaining to the limit, *v* → 0. The X-intercept of the line is *κ* = 493.33, the value of *v* for which the adsorption coefficient is zero, reflecting saturation of all the bacterial receptor sites. Except for the section comparing the model’s predictions with experimental results, the adsorption coefficient is interpreted qualitatively.

For a finite vessel, the value for *a*_*o*_ is implicitly a function of the density of the microbes, *N/* 𝒱, where is 𝒱 the volume of the vessel. For a given *N*, virus in a small vessel have more adsorption than virus in a large vessel because more virus to bacteria contacts occur in a small vessel than a large vessel during the time step. If *N* increases through time in a finite vessel, the density increases making the absorption parameter, *a*_*o*_, also increase through time. In an open infinite environment however, the local microbial density may remain relatively constant because as the microbial population spreads through space the volume of the vessel, 𝒱, effectively increases too so that *a*_*o*_ may remain relatively constant. A strong dependence of *a*_*o*_ on *N* for a fixed vessel volume would represent nonlinear feedback that may affect the phage-bacteria population dynamics. If that is important, the model can be extended by developing an expression for *a*_*o*_ following the protocol above using various values of microbial density, *N/* 𝒱.

In addition to bacterial density, *a* also depends on myriad other local environmental conditions such as media viscosity, convection from micro turbulence in the ocean or stirring in a laboratory beaker, air fans and filters, strength of the virus to bacterial binding, and temperature. Many micro processes underly the realized adsorption. The *µ* is the bottom line, so to speak—the total adsorption per time step in the local conditions integrated over all the local micro processes that bring about the adsorption. Then the *a* is the total adsorption divided by the virus to microbe ratio responsible for the adsorption.

Turning now to burst size, several choices are available for a bacterium as a function of the number of virus in it, *b*(*m*). The burst size might increase linearly with *m* so that *b*(*m*) = *b*_*o*_ *m* where *b*_*o*_ is the contribution to the overall burst made by a single viral particle (Patel and Rao 1984, Figures 1–2).

This case pertains to the absence of density dependence in burst size. Another possibility is where *b*(*m*) to be a concave increasing function of *m* (Gadagkar and Gopinathan 1980, Figures 4–5). In this case, the burst-size function might be taken as *b*(*m*) = *b*_*o*_ *m*^*α*^ where *α* is an exponent between 0 and 1, say 1/2. This case shows limited density dependence in burst size. Still a third possibility is that *b*(*m*) is a constant, *b*(*m*) = *b*_*o*_, indicating strong density dependence. This situation is described in Ellis and Delbrück (1939) who write, “The bacteria which had adsorbed several phage particles behaved as if only one of these particles was effective.” This case postulates that one infection disables all subsequent infections so that the burst is exactly the same with *m* = 1 as it is for all *m >* 1, a claim of uncertain plausibility. A final possibility is to allow *α* to be negative, say *α* = − 1, to indicate that burst size varies inversely with the multiplicity of infection, indicating extremely strong density dependence as a response to “superinfection” (Brown and Bidle 2014). All these relations between burst size and *m* may be subsumed by taking *b*(*m*) = *b*_0_ *m*^*α*^ with *α* as 1, 1/2, 0, or -1. The cases of *α* = 1 and *α* = 0 were presented in Roughgarden (2024) and results for other *α* were noted, but not shown.

Here, the burst size as a function of the multiplicity of infection is taken as

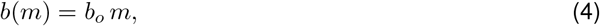

where *b*_*o*_ is the burst per virus infecting the bacterium. Even if *b*(*m*) is not linear throughout the range of possible *m*, the linear form can be viewed as a Taylor approximation in the limit as *m* → 0. This approximation is appropriate because the subsequent analysis focusses on the conditions for the virus to increase when rare into a bacterial population that is growing in its log phase—in such a circumstance *m* is small.

With *a*(*v*(*t*)) from Eq. 3, the equation for *N* (*t* + 1) from Eq. 1, becomes explicitly

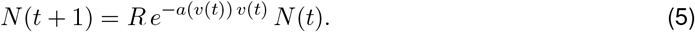

The sum in the equation for *V* (*t* + 1) from Eq. 2 can be found using properties of the Poisson distribution combined with *b*(*m*) from Eq. 4, yielding

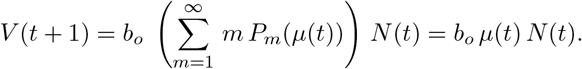

Because *µ*(*t*) = *a*(*v*(*t*)) *v*(*t*), and *v*(*t*) ≡ *V* (*t*)*/N* (*t*), the equation above becomes

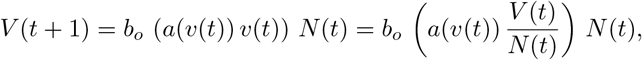

which reduces to

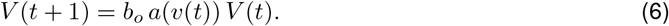

Both the equations for *N* (*t* + 1) and *V* (*t* + 1) involve the virus to microbe ratio, *v*(*t*). The dynamical equation for their ratio is found by dividing Eq. 6 by Eq. 5 yielding

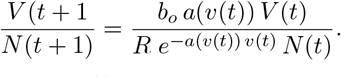

Then identifying 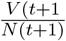 with *v*(*t* + 1) and 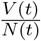 with *v*(*t*) yields,

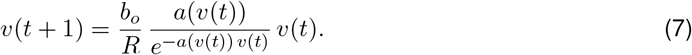

Equation 7 is a dynamical equation in one variable, *v*(*t*), identical to Eq. 8 in Roughgarden (2024). The *v*(*t*) may approach a stable virus to microbe ratio, 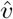, at which both the virus and bacteria grow geometrically with the same factor each time step. Moreover, even if 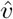 exists and is stable, the virus-bacteria pair may nonetheless go extinct because, in Eq. 5, *R* × the factor, 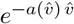, may be less than one, even though *R* is greater than 1. This factor, called the “infection discount” in Roughgarden (2024), indicates the degree to which the bacterial population growth factor is discounted as a result of the morality caused by viral infections. The infection discount is generally less than 1 but approaches 1 as 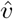 approaches 0, indicating that a low virus to microbe ratio has little effect on the net bacterial growth. In contrast, a high virus to microbe ratio implies that many microbes are being infected and killed. This strongly discounts the intrinsic microbial geometric growth factor, *R*, so that the net geometric growth factor can be less than one even though the geometric growth factor for the uninfected microbes is greater than one.

Unlike Roughgarden (2024), this paper does not focus on the stable virus to microbe ratio, 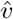. Instead, the focus is on the condition for increase when rare for a virus, that is, the condition when a virus can invade and infect a population of uninfected microbes. What traits does a virus need, and specifically what lysis time and burst size does it need, to invade faster than other viruses with other lysis times and burst sizes?

So, in the limit, *v*(*t*) → 0, the dynamical equation in Eq. 7, upon suppressing the subscripts, reduces to

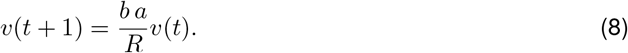

The virus population can infect the bacterial population if it can increase when rare, *ie*., if *v*(*t*) ≈ 0, then *v*(*t* + 1) *> v*(*t*). Inspection of Eq. 8 shows that if

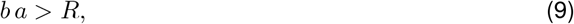

then the virus can spread into a population of increasing bacteria. Alternatively, for a virulent virus, if *b a/R <* 1, the virus goes extinct because the virus cannot keep up with bacterial population growth. But for a temperate virus, if *b a/R <* 1, the virus switches to a prophage because its population grows faster as part of the bacterial genome than by bursting new virions each time step. This criterion was introduced in Roughgarden (2024).

#### Optimal Balance Between Burst Size and Lysis Time

This section applies the phage-bacteria population-dynamic model of the previous section to the tradeoff between burst size and lysis time.

Wang (2006) experimentally determined a functional relation between burst per infecting virus and lysis time as

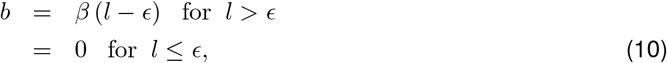

where *l* is the lysis time, *ϵ* is the eclipse, which is the time needed for the viral replication to commence, and *β* is the phage production rate (also called the phage maturation rate) once viral production has begun. The burst per virus increases linearly with the lysis time period after the eclipse has transpired. (These symbols and all the others used in the paper are listed in Table 1.)

**Table 1:** Symbols.

| Symbol | Description | Units |
| --- | --- | --- |
| $V$ | Number free virus particles (virions) | Organisms |
| $N$ | Number of microbes (bacteria) | Organisms |
| $v$ | Virus to microbe ratio, VMR, | None |
| $R$ | Geometric factor of increase per time step for microbes | None |
| $\rho$ | Geometric factor of increase per minute for microbes | None |
| $\mu$ | Poisson density, average total adsorption per time step | Organisms |
| $P_m(\mu)$ | Probability microbe colonized by $m$ virus | None |
| $a(v)$ | Adsorption function, total adsorption per VMR | Organisms |
| $a_o$ | Maximal total adsorption per VMR when VMR is low | Organisms |
| $\kappa$ | Saturation VMR | None |
| $b(m)$ | Burst per microbe as function of multiplicity of infection | Organisms |
| $b_o$ | Burst per infecting virus | Organisms |
| $a$ | $a_o$ with subscript suppressed | Organisms |
| $b$ | $b_o$ with subscript suppressed | Organisms |
| $l$ | Lysis time (latency), same as time step | Minutes |
| $L$ | Large time interval to include multiple time steps | Minutes |
| $\epsilon$ | Eclipse period, time before virus production starts | Minutes |
| $\beta$ | Virus production rate once eclipse expires | Organisms per minute |
| $W_L$ | Relative fitness of lytic to lysogenic phage over interval $L$ | None |
| $Y(l)$ | Expression whose root is optimal lysis time | None |
| $\hat{l}$ | Optimal lysis time | Minutes |
| $\hat{b}$ | Burst per infecting virus with optimal lysis time | Organisms |
| $l_c(L)$ | Critical lysis time for lytic = lysogenic fitness over interval $L$, | Minutes |
| $W_{Lo}$ | $W_L$ assuming $\epsilon$ is zero | None |
| $\hat{l}_o$ | Optimal lysis time assuming $\epsilon$ is zero | Minutes |
| $\hat{b}_o$ | Optimal burst size assuming $\epsilon$ is zero | Organisms |
| $\widehat{W_{Lo}}$ | Optimal $W_L$ assuming $\epsilon$ is zero | None |

The geometric growth factor of uninfected microbes, *R*, in Eq. 1 refers to microbial population growth over a time step. Because the lysis time, and hence the time step, can vary it is useful to refer the geometric factor to units of a minute, rather than to the time step itself. Hence, define *ρ* as the uninfected microbe geometric growth factor per minute. Then the growth factor for a time step of length, *l*, is

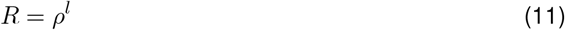

Because the time step in the model, Eq. 8, is the lysis time, *l*, to investigate how the time step itself evolves, a maximum time step, *L*, is stipulated that is longer than the largest feasible lysis time. Then the dynamical equation is iterated for the number of *l* that fit within *L*, given the eclipse. For example, suppose the eclipse is 0 and *L* is 100. If *l* is also 100, then only one iteration takes place within *L*, whereas if *l* is 1, then 100 iterations fit within *L*, and if *l* is 25, then four iterations fit within *L*, and so forth. Thus, for a given *l*, the *v* after the number of iterations that fit within *L* is

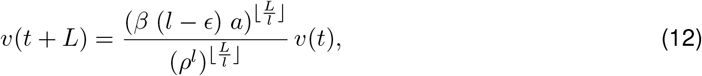

for *l* = (*ϵ*+1, *ϵ*+2 … *L*) where 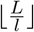 indicates the integer part of 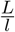. This is the number of iterations of viral reproduction with lysis time, *l*, that fit into *L*.

The numerator of Eq. 12 is the production of virions from a lytic virus over the number of time steps of size, *l*, that fit into the interval, *L*. The denominator of Eq. 12 is the bacterial production over the number of time steps of size, *l*, that fit into the interval, *L*. This is also the production of a temperate virus residing as a prophage in the bacterial chromosome. The geometric growth factor for a microbe over a single time step of length *l* is *ρ*^*l*^. The *l* in the exponents within the denominator cancel leaving,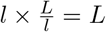

Further analysis is aided by taking a continuous approximation in Eq. 12, as though a fractional iteration can fit into *L*,

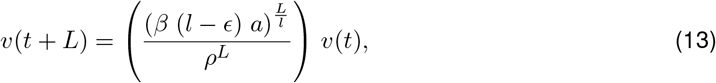

for *l > ϵ*. The expression in large parentheses in Eq. 12 is the relative fitness of a lytic phage relative to the prophage residing in the bacteria over the period, *L*, as a function of *l*,

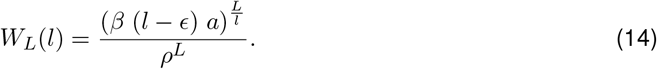

The optimal lysis time, 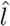, is found by maximizing *W*_*L*_(*l*) with respect to *l*. The corresponding optimal burst per infecting microbe, 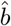, is found from Eq. 8, evaluated at 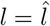

To find the optimal lysis time, differentiate *W*_*L*_ with respect to *l*, yielding

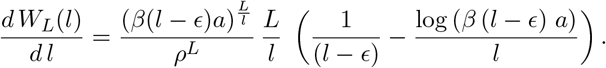

This equation contains the expression in large parentheses defined as

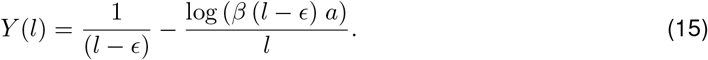

The root, *Y* (*l*) = 0, of this expression is the optimal lysis time, 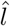. The *Y* (*l*) is independent of both *L* and *ρ*, and therefore, so is 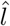

The critical lysis time, *l*_*c*_(*L*), at which being lytic becomes a better strategy than being a prophage over the period, *L*, is found by setting

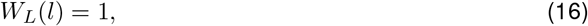

and solving for *l*. If the optimal lysis time, 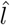, is less than *l*_*c*_(*L*), the virus is better off not lysing at all during the period, *L*, and instead incorporating as a prophage into the bacterial chromosome. Alternatively, if 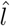 *l*, is greater than *l*_*c*_(*L*), the virus is better off infecting and lysing the bacteria during the period, *L*, while doing so with the optimal lysis time. The optimal lysis time produces a corresponding optimal burst per infecting virus found from Eq. 10 evaluated at 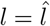

### Predictions

#### Comparison of Model with Laboratory Data

Table 2 gives the parameters for an illustration. The *β* and *ϵ* in units of phage/min and min, respectively, are derived from isogenic *λ*-phages with different lysis times from Wang (2006). The bacterial geometric factor of increase, *R*, from Wang (2006) for *Escherichia coli* K-12, is given as 2.04/hr—this converts to the value of *ρ* in the table in minutes. In the absence of an experiment to measure the adsorption parameter, *a*, a nominal value of 0.1 is used corresponding to the worked example provided earlier. The adsorption parameter varies considerably depending on environmental conditions affecting the accessibility of the bacteria to the virus and the number of binding sites per bacterium varies with nutrient conditions (Schwartz 1976, Philipson 1983). The large time interval, *L*, is taken arbitrarily as 100 minutes, a value comfortably larger than the optimal lysis time.

**Table 2:**
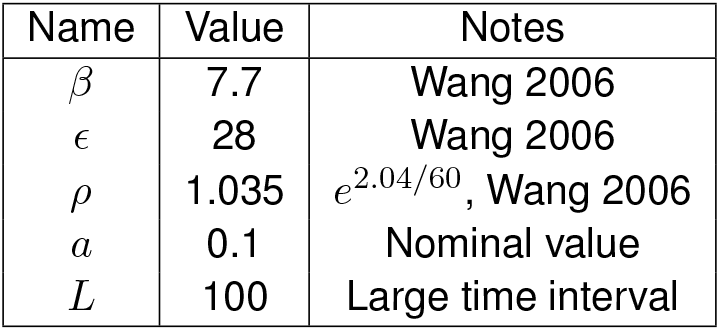
Parameters.

| Name | Value | Notes |
| --- | --- | --- |
| $\beta$ | 7.7 | Wang 2006 |
| $\epsilon$ | 28 | Wang 2006 |
| $\rho$ | 1.035 | $e^{2.04/60}$, Wang 2006 |
| $a$ | 0.1 | Nominal value |
| $L$ | 100 | Large time interval |

To find the optimal lysis time, using the NSolve[] function in Mathematica with *Y* (*l*) in Eq. 15, together with the parameters from Table 2, returns the value,

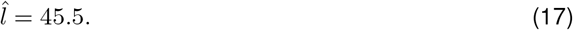

independent of *L* and *ρ*. The corresponding optimal burst per infecting virus from Eq. 10 is

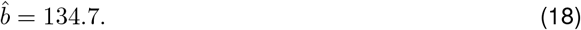

and the corresponding maximal relative fitness over 100 minutes from Eq. 14 is

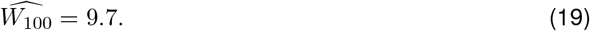

and the critical lysis time at which being lytic becomes a better strategy than being a prophage over 100 minutes works out to be

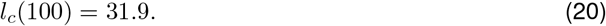

Figure 2 shows the fitness of a lytic phage relative to a prophage after a period of 100 minutes as a function of the lysis time, *l*, based on the parameters in Table 2. Relative lytic to prophage fitness of 1 is shown as horizontal line. If *l* is greater than 31.9, then the lytic phase has a higher fitness than the lysogenic phase during the 100 minute interval. If *l* is between *ϵ* and *l*_*c*_(100), *i*.*e*. between 28 and 31.9 minutes, then the phage cannot coexist during the interval with the bacteria in its lytic phase—the bacteria population grows faster than the viral population, leaving it behind, so to speak. Instead, the phage must lysogenize—incorporate into the bacterial genome to acquire the population growth rate of the bacteria.

**Figure 2:**
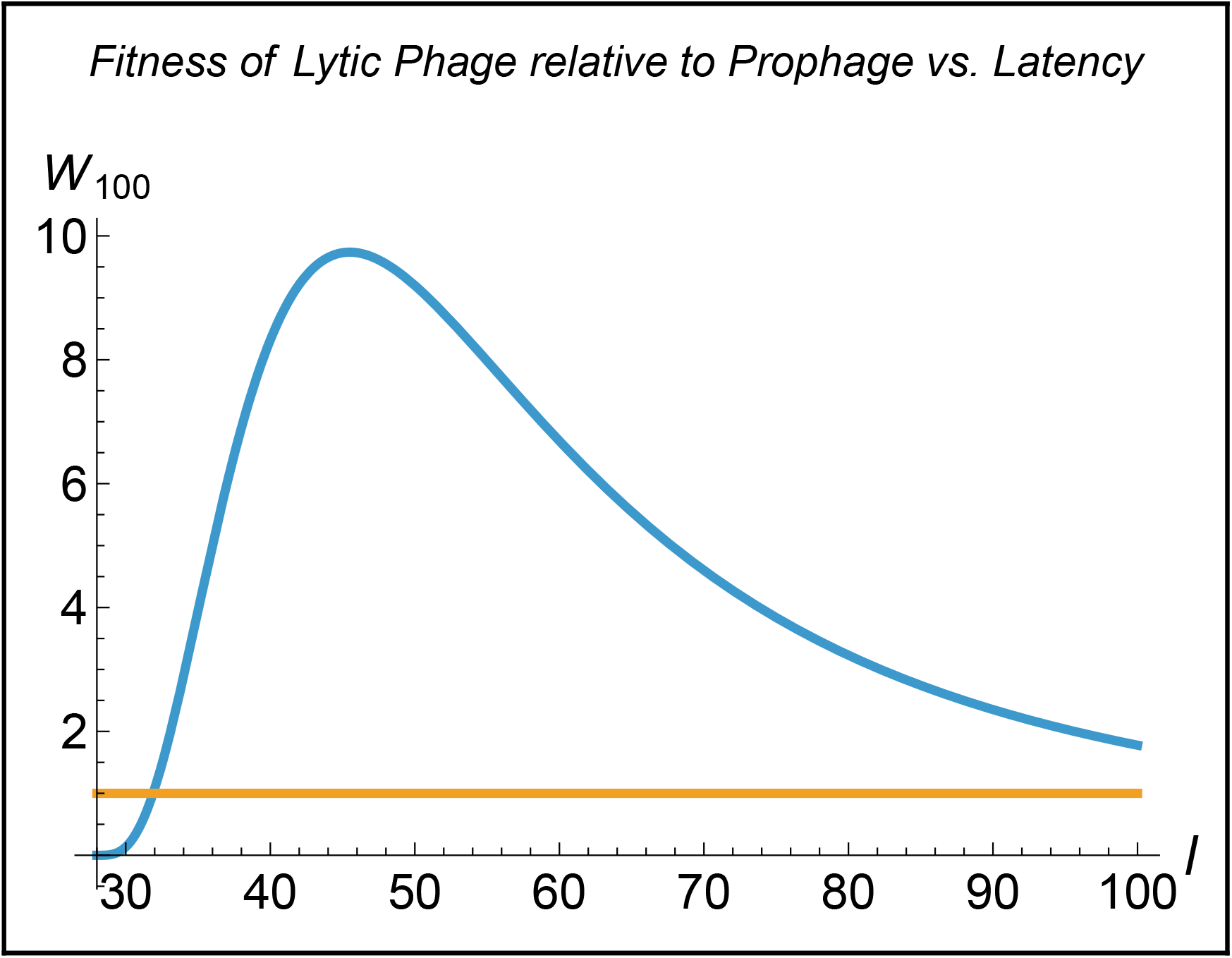
Fitness of lytic phage relative to prophage after 100 minutes, W_100_, as a function of the lysis time, l, is a humped-shaped curve starting from the eclipse period of 28 minutes. The optimal lysis time occurs at 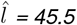 which yields a maximum ratio of lytic to prophage fitness, 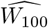, of 9.7 over the 100 minute interval. Relative lytic to prophage fitness of 1 is shown as horizontal line. The hump-shaped curve for W_100_ intersects the line at l_c_(100) = 31.9. Thus, the fitness of a lytic phage exceeds that of a prophage during the 100 minute interval when the lysis time exceeds 31.9. If the lysis time of the phage is between 28 and 31.9, the phage cannot coexist with the bacteria during the 100 minute interval unless it becomes a prophage.

Wang (2006) measured the fitness of 11 strains of *λ*-phage whose latencies varied from 28.3 to 63.0 minutes. The study found that the strain whose lysis time was 51.0 minutes had the highest fitness, a value quite close to that predicted in the model here, 45.5, based on the parameters in Table 2. The burst per infecting virus for the strain with the highest fitness was 170.0, also fairly close to the value predicted here, 134.7. This initial degree of agreement between theory and experiment is coincidental because the nominal value of the adsorption parameter, *a*, in Table 2 was not experimentally determined.

Instead, suppose *a* is lowered to 0.052. Then 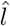 works out to be 50.0 and 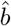to 176.9, almost exactly the same as Wang(2006) found experimentally. Thus, lowering *a* from the nominal value of 0.100 to 0.052, while keeping the other parameters from Table 2 the same, brings the predicted optimal lysis time and burst per infecting virus into close agreement with the experimental results. The next section considers further implications of variation in *a*.

#### Interventions that Impact Adsorption

Figure 3 shows the impact of varying adsorption on phage evolution. The figure starts with *a* at a low value for which the phage exists as a prophage. Then increasing *a* induces the phage to excise from the bacterial chromosome when the ratio of lytic to prophage fitness equals 1. Thereafter, as *a* increases even more, the optimal latency and bust size progressively decrease while the relative fitness of the lytic to prophage progressively rises.

**Figure 3:**
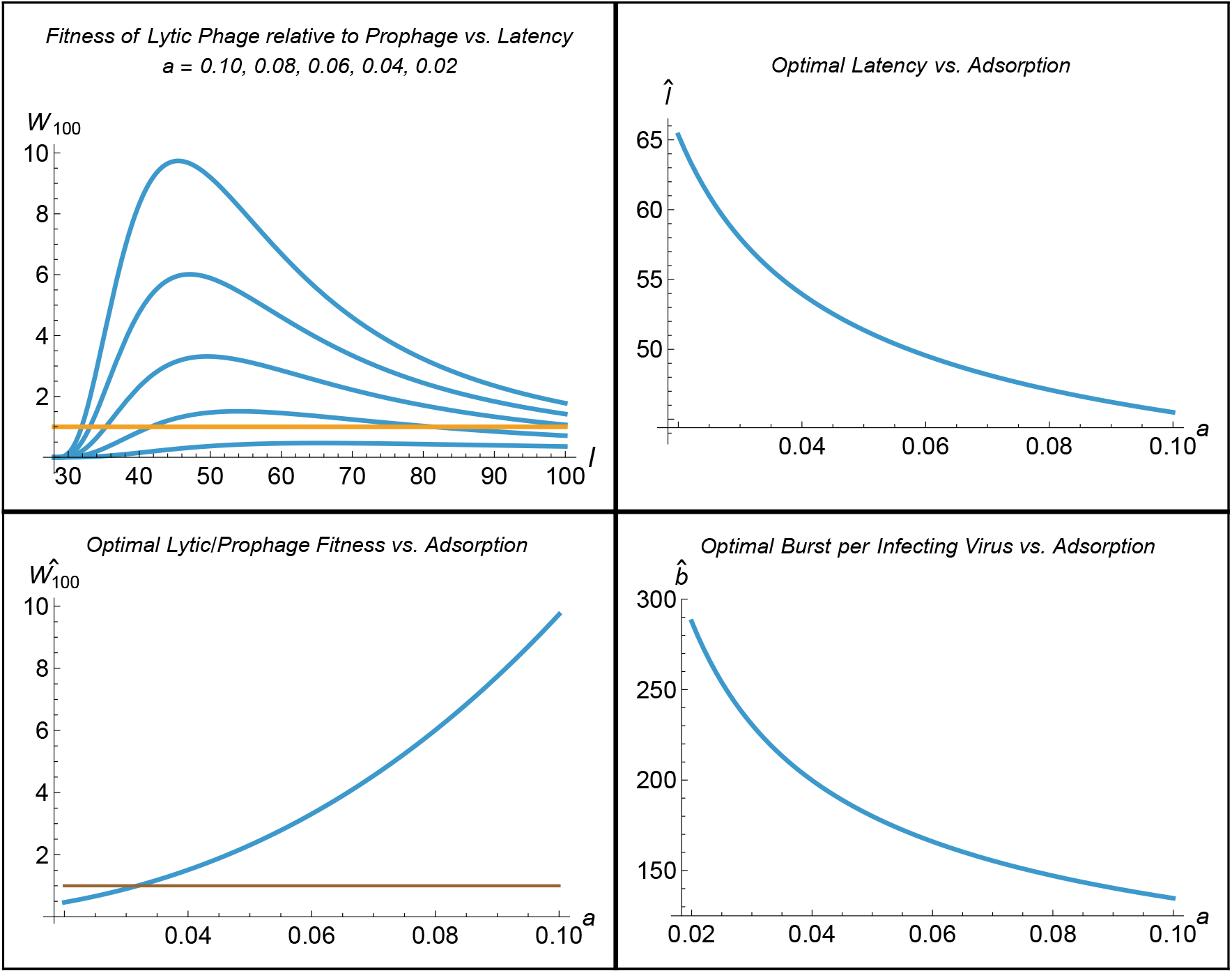
Impact of varying adsorption on phage evolution. Top left: Fitness of the lytic phase vs. lysis time for several values of the adsorption coefficient, a. The top curve is for a = 0.10 and is identical to that shown in Figure 2. The curves below the top are successively for a = 0.08, 0.06, 0.04, and 0.02. The horizontal line is a relative fitness of 1 where the fitness of a lytic phage equals that of a prophage. As a decreases, the peaks of the curves for the lytic phase progressively become lower and shift to the right. Top right: Optimal lysis time vs. the adsorption parameter, a. As a increases from 0 to 0.10, the optimal lysis time decreases. Bottom right: Optimal burst per infecting virus vs. the adsorption parameter, a. As a increases from 0 to 0.10, the optimal burst size also decreases, reflecting the decreasing lysis time. Bottom left: Optimal fitness of a lytic phage relative to the prophage vs. the adsorption parameter, a. The horizontal line is a relative fitness of 1 where the fitness of a lytic phage equals that of a prophage. As a increases from 0 to 0.10, the optimal lytic/prophage fitness ratio starts below 1 and increases, intersecting the horizontal line at a = 0.032, at which point the prophage becomes induced to excise from the bacterial chromosome and become lytic.

Many interventions to control phages, and viruses more generally, involve techniques such as air or water filters, or controlling humidity that lower the adsorption of phage by bacteria (*cf*. Sobsey *et al*. 2008, Clokie and Kropinski 2009, Wommack *et al*. 2009, Moriyama *et al*. 2020, Curtius *et al*. 2021, Van Charante *et al*. 2021, Beltrán *et al*. 2023). Overall, the model predicts that interventions lowering *a* cause the evolution of an increased lysis time and correspondingly higher burst size until the intervention is sufficient either to drive a virulent phage extinct, or for temperate phage, to drive the phage from its lytic phase into its lysogenic phase.

#### Eco-geographic Patterns

Prevalence of the lysogenic life cycle relative to the lytic life cycle increases in conditions that promote bacterial productivity. For example, overfishing and eutrophication on coral reefs causes an increase in ungrazed fleshy algae. These release dissolved organic carbon that promotes bacterial productivity, leading to a condition called microbialization. Knowles *et al*. (2016) and Silveira *et al*. (2021) show that lysogeny increases with microbialization on coral reefs. Another example involves the guts of mammals. Here concentrated nutrients promote bacterial productivity, resulting in gut bacteria that are enriched with prophages, (López-Leal *et al*. 2022). Generally speaking, increasing bacterial productivity relative to virion productivity increases the likelihood of the lysogenic lifecycle relative to lytic lifecycle because reproduction as a prophage becomes better than reproduction through lysis (Roughgarden 2024).

Not as many eco-geographic patterns are known for burst size and latency time as for the lysogenic to lytic ratio. However, in one study, Parada *et al*. (2006, Fig. 5) showed a positive relationship between bacterial productivity and viral burst size in the North Sea. In terms of the model, high nutrient conditions increase both *ρ*, the bacterial geometric growth factor and perhaps *β*, the number of new virions produced per infecting virus per minute of latency. High nutrient conditions might affect also *a*, the absorption coefficient depending on whether high nutrients increase or decrease bacterial density.

The eclipse period may a quarter to a half of the overall latency, being about 10 minutes for the phage T7 whose latency is 40 minutes at 30^◦^ (Endy *et al*. 1997, Fig. 6) and 28 minutes for *λ* phage with an optimal latency of 51 minutes, as discussed earlier (Wang 2006, *cf*. Table 2). Still, qualitative insight might be gained by letting *ϵ* be zero. Accordingly, Eq. 14 becomes

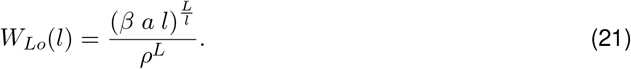

By differentiating *W*_*Lo*_(*l*) with respect to *l*, setting the result equal to zero and solving for *l*, yields an explicit result for the optimal latency

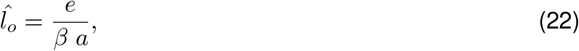

where *e* is Euler’s Number, the base of the natural logarithms, and not the eclipse, *ϵ*. Thus, the optimal latency for the virus varies inversely with how good the environment is for the lytic phase. The more favorable the environment is for the lytic phase, (*β a*), the shorter the optimal latency. That is, the better the environment is for the lytic phase, the faster the virus goes around its life cycle. Specifically, Figure 4 (Left) shows 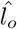 as a function of (*β a*) from Eq. 22.

**Figure 4:**
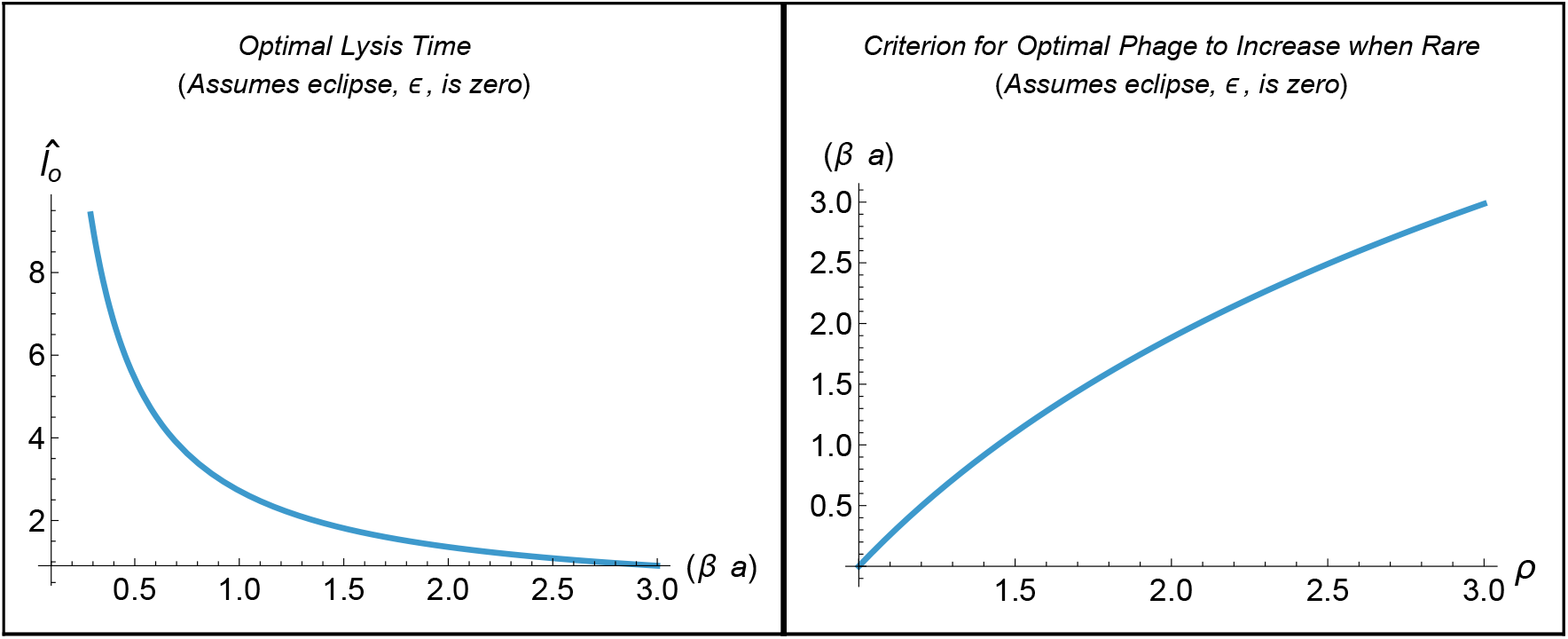
Left: The optimal lysis time 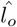 as a function of the quantity (β a), where β is the rate of virion production per minute of lysis time and a is the adsorption parameter, from Eq. 22. Right: Graph of criterion for increase when rare for an optimal lytic phage as a function of the bacterial geometric growth factor, ρ, from Eq. 27. If (β a) is greater than the curve of (e log[ρ]) then the virus can increase when rare, otherwise it is excluded from the system if virulent, or becomes lysogenic if temperate. If the lytic virus can increase when rare, the corresponding optimal lysis time can be read off the graph on the left.

The burst per infecting virus realized during the optimal latency, from Eq. 10 with *ϵ* = 0, is

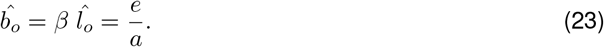

The optimal burst per infecting virus is independent of *β*. To see why, suppose productive environmental conditions increase the speed at which new virions are produced, *β*. This leads to a shorter optimal Latency,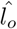. Then the higher speed of virion production times the shorter latency cancel out leaving the burst per infecting virus unchanged. However, the optimal burst per infecting virus does vary inversely with *a*. A lower bacterial density leading to a lower *a* causes a higher burst per infecting virus according to Eq. 23.

The Parada *et al*. (2006) observations in which both bacterial productivity and viral burst size increase together across various sites in the North Sea can be reconciled with this model if both (1) higher nutrients increase *ρ*, as was observed, and (2) higher nutrients also decrease bacterial density, thereby lowering *a*, which in turn leads to higher burst per infecting virus according to Eq. 23. Also according to Eq. 23, any effect of nutrients on *β* is irrelevant. This ecological situation is where high nutrients cause high productivity and turnover associated with a low standing crop of bacteria, as is characteristic of grazed ecosystems (*eg*. Odum and Barrett 2005).

The virion productivity per the time step equalling the optimal latency is invariant because 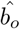 depends inversely on *a*, in Eq. 23. The *a* therefore cancels out in

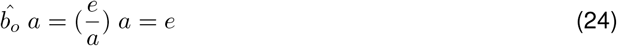

To see why, suppose the adsorption, *a*, increases. Then the optimal latency decreases. Therefore, the corresponding burst per infecting virus decreases too. The decreased burst per infecting virus then cancels out the increase in adsorption leaving the virion productivity per the time step corresponding to the optimal latency unchanged. So, as *a* and *β* are varied across environmental gradients, the optimal latency does vary according to Eq. 22. However, the virion productivity resulting from those varying latencies remains constant for the corresponding time steps.

The relative advantage of the lytic to lysogenic lifecycle can to be computed over a long period, *L*. Multiple iterations of the single-step model occur within this period that can be compared with bacterial population growth over the same period. The optimal virion productivity over *L* relative to the bacterial productivity over *L* is found by substituting Eq. 22 into Eq. 21 leading to

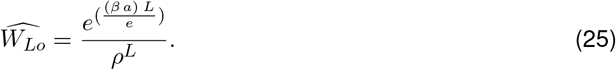

The 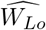 increases with (*β a*) and decreases with *ρ*. To see the long-term relative advantage of the optimal lytic lifecycle to the lysogenic lifecycle, take the limit as *L* tends to infinity, yielding

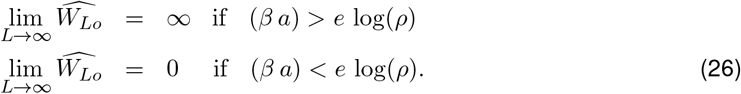

Thus, the optimal virus, *i*.*e*., one with the optimal balance between latency and burst size, increases when rare over the long term if

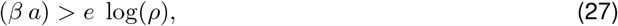

otherwise it goes extinct or becomes lysogenic. That is, for an optimal virus, its productivity as a virion must exceed its productivity as a prophage to be virulent according to Eq. 27.

Figure 4 (Right) shows the criterion for increase when rare for an optimal lytic phage as a function of the bacterial growth factor, assuming the eclipse is zero, from Eq. 27. If (*β a*) on the vertical axis is greater than the curve for (*e* log(*ρ*)) then the virus can increase when rare, otherwise it is excluded from the system if virulent, or becomes lysogenic if temperate.

To pull these results together, consider a gradient in environmental factors along which the lytic productivity, (*β a*), increases. Assume too that the eclipse, *ϵ*, is small. Then (1) the optimal lysis time shortens along the gradient, implying that the lytic life cycle goes around faster along the gradient, according to Eq. 22; (2) The burst per infecting virus does not vary along the gradient, according to Eq. 23, unless *a* changes because of bacterial density change along the gradient; and (3) the virion productivity realized during the time step corresponding to each point along the gradient remains constant, according to Eq. 24. If the bacterial productivity, *ρ*, also increases along this gradient, then the condition in Eq. 27 can flip. At the flipping point on the gradient, the virus either goes extinct or if temperate, switches to lysogeny.

## Discussion

This article provides a mathematical model for the evolutionarily optimal balance between lysis time and burst size. Wang (2006) found the optimal lysis time with strains of *λ*-phage infecting *E. coli* to be 51 minutes. Kannoly *et al*. (2022) also measured strains of *λ*-phage infecting *E. coli* and found an optimal lysis time of 80 minutes. The value of the optimal lysis time differed between these studies because the environmental conditions in the experiments differed. Given the reality of an optimal lysis time that depends on environmental conditions, the issue is how to explain it.

This article explains the optimal lysis time by finding the optimal balance between rapid reproduction with a short lysis time at the cost of a low burst size *vs*. slow reproduction with a long lysis time and the benefit of a large burst size. The optimal balance between these two considerations is derived using a dynamical equation in discrete time for the virus to microbe ratio (VMR) during the log phase of phage/bacteria population growth (Roughgarden 2024).

Previous models for the optimal balance use a system of delay-differential equations for the dynamics of uninfected bacteria, infected bacteria and free phage in the stationary phase of virus-bacterial population growth (*eg*. Bull 2006). The system of equations is an extension of the original delay-differential equation model introduced by Campbell (1961).

In the present context, the relevant distinction between this paper and previous papers is the intuition behind the optimal lysis time. Eq. 4 in Bull (2006) gives a formula for the optimal lysis time, (called *L*). When the phage is at population-dynamic equilibrium (called 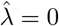) and when the eclipse (called *E*) is zero, then the optimal lysis time is *L* = 1*/d*, where *d* is the death rate of bacterial cells. This prediction from Eq. 4 in Bull (2006) for the optimal lysis time is completely different from Eq. 22 in this paper.

The intuition behind Eq. 4 of Bull (2006) for the optimal lysis time is that a high bacterial death rate implies a short life span for the bacteria, leaving only a short time for an infecting virus to make new virions. Thus, the tradeoff is between the time needed to produce a burst and the time available within a bacterial cell to do it. This explanation for an optimal balance between burst size and the time to manufacture the burst is conceptually different from that offered by this paper’s model that focusses on the balance between rapid reproduction with low burst *vs*. slow reproduction with high burst.

Using the ecological terminology of *r*-selection and *K*-selection (MacArthur and Wilson 1967, Roughgarden 1971), the present model views virus as *r* selected because the virus and bacteria are growing exponentially and the Bull (2006) model views virus as *K*-selected because the virus and bacteria are at equilibrium.

Another conceptual difference between Eq. 4 of Bull (2006) for the optimal lysis time *vs*. Eq. 22 here is that, colloquially speaking, in the former the virus is trying to synchronize its lysis time with the bacterial dynamics, whereas here, the virus is trying to find the best balance between early and late lysis irrespective of what the bacteria are doing. Then once the lytic virus is doing the best it can, it compares what it achieves by remaining lytic with what it achieves by lysogenizing and decides to remain lytic or to switch to lysogeny according to which yields the higher fitness.

Grounds for preferring this paper’s model over previous models, in addition to its simplicity and tractability, rest on which environmental situation is more common—log phase growth or stationary growth for the phage-bacteria pair. Previous models are directed to experimental situations such as a chemostat where a stationary phase can be realized. But in nature it is hard to imagine phage and bacteria in equilibrium. In nature the populations are invariably either recovering from or being decimated by environmental events such as storms *etc*. Natural populations of phage and bacteria are likely continually in a bust-boom regime where a log-phase model is preferable to a stationary-phase model.

The model of this paper predicts the evolutionary impact of interventions to control phage and predicts possible eco-geographic patterns of latency and burst size along environmental gradients. The model predicts that interventions lowering phage adsorption cause the evolution of an increased lysis time and correspondingly higher burst size until the intervention is sufficient either to drive a virulent phage extinct, or for temperate phage, to drive the phage from its lytic phase into its lysogenic phase. The model predicts that along a gradient in which the viral productivity increases, the optimal lysis time shortens and the lytic life cycle goes around faster.

All in all, this paper shows that the life-history traits of virus are not arbitrary, but vary in predictable ways associated with different environments.

## Acknowledgments

The author thanks Cynthia Silveira of the University of Miami together with Forest Rohwer and members of the San Diego State University (SDSU) BioMath Group for discussion about this paper. This article is Contribution #4 from a project, The Theory of Holobiont Evolution, funded by the Gordon and Betty Moore Foundation through Grant GBMF10000 to the University of Hawaii, J. Roughgarden, PI.

